# Strict Early Restraint of Anterior Tibial Translation Favors Spontaneous Anterior Cruciate Ligament Healing: Implications for Non-Operative Management from a Murine Rupture Model

**DOI:** 10.1101/2024.05.28.596036

**Authors:** Riku Saito, Kazuki Nakayama, Yuna Usami, Saaya Enomoto, Koyo Nogi, Takanori Kokubun

## Abstract

**Background:** Anterior cruciate ligament injury is a sports-related injury traditionally considered to have limited spontaneous healing potential. Recent evidence suggests that healing can occur under appropriate mechanical conditions; however, non-operative protocols differ in the anterior tibial translation they permit, and the magnitude compatible with healing remains unclear. This study examined how the magnitude of anterior tibial translation restraint affects spontaneous ligament healing and tissue maturation.

**Methods:** Male and female C57BL/6 mice (n = 142) underwent non-invasive anterior cruciate ligament rupture and anterior tibial translation was restrained strictly or mildly with extra-articular sutures, or left unrestrained. Knees were assessed at 0, 4, and 8 weeks after injury. Outcomes were histological healing, collagen organization, anterior tibial translation, and failure load.

**Findings:** Strict restraint reduced anterior tibial translation to levels not significantly different from intact knees immediately after injury, although this effect diminished by 4 weeks. After adjustment for sex and time point, strict restraint was associated with higher odds of healing than mild restraint (adjusted odds ratio, 4.29; *p* = 0.019), whereas no knees healed without restraint. Among healed ligaments, mild restraint did not improve collagen organization or failure load, and both remained inferior to intact ligaments.

**Interpretation:** Strict restraint of anterior tibial translation during the early phase was more favorable for re-establishing ligament continuity than permitting mild translation, which did not benefit tissue maturation. For non-operative management, early protocols may need to prioritize restraint of anterior tibial translation, deferring loading until continuity is established; clinical studies should define this timing.

**Highlight:**

- Restraint magnitude altered anterior cruciate ligament spontaneous healing outcomes.
- Strict early restraint favored spontaneous healing after complete ligament rupture.
- Mild anterior tibial translation did not improve tissue maturation or strength.
- Healed ligaments remained structurally and mechanically inferior to intact ligaments.

## 1 Introduction

Anterior cruciate ligament (ACL) injury is one of the most common sports-related injuries (1), with more than 100,000 cases reported annually in the United States, imposing a substantial medical and socioeconomic burden (2). The ACL has been considered to have limited intrinsic healing capacity because of its relatively poor vascular and cellular responses compared with the medial collateral ligament (MCL) (3–6). Consequently, ACL reconstruction has been widely performed as the standard treatment to restore knee stability and function after ACL rupture. However, concerns regarding growth plate injury in skeletally immature patients (7) and postoperative osteoarthritis progression in older patients (8) have renewed interest in strategies that promote intrinsic ACL healing.

Recent clinical and preclinical studies have increasingly reported the potential for spontaneous ACL healing (9–15). Ihara et al. first reported that controlling abnormal mechanical forces with functional bracing could facilitate ACL healing (9). More recently, Filbay et al. reported evidence of ACL healing in 90% of patients treated with the Cross Bracing Protocol, further highlighting the importance of controlling the mechanical environment during the early phase after ACL injury (15). Because the ACL is the primary restraint to anterior tibial translation (ATT), its rupture increases ATT, which may separate the ligament remnants and expose the healing tissue to excessive strain (10,16). However, the optimal mechanical conditions for promoting successful ACL healing remain unclear. Current clinical approaches vary from functional bracing intended primarily to control ATT to protocols involving more rigid restriction of knee motion. These approaches permit different magnitudes of ATT, but the magnitude compatible with healing cannot be systematically controlled or quantified in patients, which necessitates an animal model in which ATT restraint can be graded. In our previous animal study, rigid ATT restraint facilitated spontaneous healing of completely ruptured ACLs; however, the healed ligaments remained structurally and mechanically immature (10). Appropriate mechanical loading can promote tissue repair and maturation in ligament and tendon healing (17–21), whereas insufficient loading may impair tissue maturation (22,23). Achieving both successful ACL healing and subsequent tissue maturation may therefore require a balance between sufficient ATT restraint to maintain tissue continuity and mechanical loading to promote maturation.

Therefore, the purpose of this study was to determine how the magnitude of ATT restraint influences spontaneous ACL healing, subsequent tissue maturation, and the knee stability and mechanical properties of the healed ACL. To address this question, we developed two controlled anterior tibial translation (CATT) models using nylon sutures of different diameters to provide different levels of ATT restraint. We hypothesized that mild residual ATT, allowing increased mechanical loading of the healing ACL, would permit spontaneous healing while enhancing tissue maturation and mechanical recovery compared with strict restraint.

## 2 Methods

### 2.1 Experimental Design

A total of 142 C57BL/6 mice (71 males and 71 females, 8–9 weeks old) underwent non-invasive rupture of the left ACL and were randomly assigned to the Tight-CATT (3-0 nylon suture, n = 57), Loose-CATT (4-0 nylon suture, n = 55), or ACL-Rupture (ACL-R; uncontrolled ATT, n = 30) group (Fig. 1A). Allocation was performed using simple randomization using a computer-generated random sequence, and the sample size was determined with reference to previous studies (10,12,24). Investigators performing ATT measurements, histological classification, and image analyses were blinded to group allocation. The contralateral right hindlimb served as the intact control. Mice were housed under a 12-h light/dark cycle with unrestricted cage activity and free access to food and water. Six mice were excluded because of rupture-related fractures or experimental abnormalities (Tight-CATT, n = 3; Loose-CATT, n = 3), leaving 136 mice for analysis. Mice were sacrificed immediately after injury (n = 30), at 4 weeks (n = 52), or at 8 weeks (n = 54) (Fig. 1B). Analyses of collagen organization, COL1A1 localization, knee stability, and biomechanical properties were performed only on healed ACL samples.

**Fig. 1.**
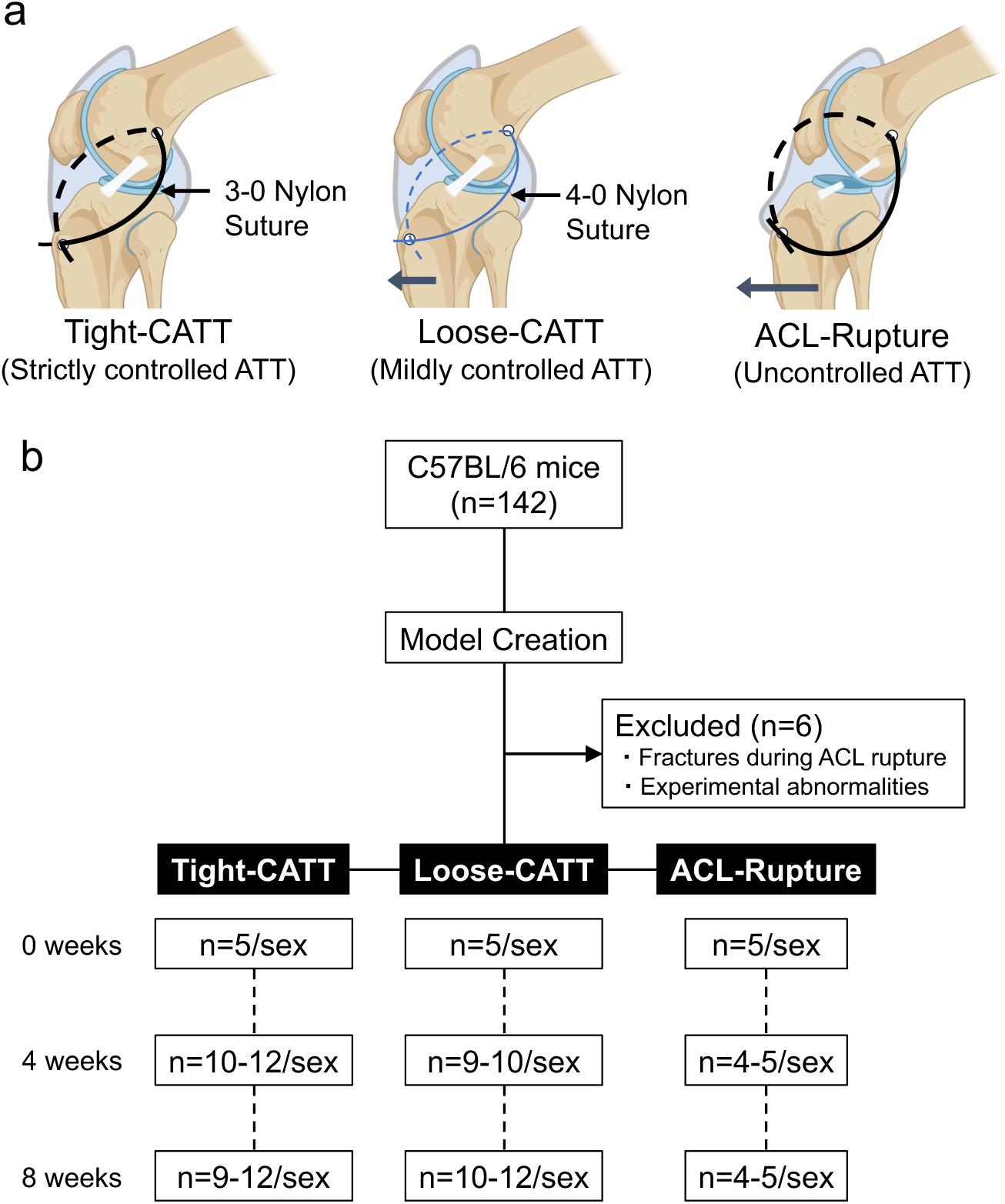
Experimental design. (A) Following noninvasive ACL rupture, mice were assigned to Tight-CATT (larger-diameter 3-0 nylon suture), Loose-CATT (smaller-diameter 4-0 nylon suture), or ACL-Rupture (ACL-R; no ATT restraint). (B) Study flow diagram. After exclusion of six mice, knee tissues were harvested immediately and at 4 and 8 weeks; numbers per sex, group, and time point are shown.

### 2.2 Procedure for Creating the Model

All mice were anesthetized with medetomidine (0.375 mg/kg), midazolam (2.0 mg/kg), and butorphanol (2.5 mg/kg), and the ACL was ruptured non-invasively as previously described (25). The hindlimb was fixed with the knee at 90°, and ACL rupture was induced by pushing the femur posteriorly against the tibia. Bone tunnels were then created at the femoral condyle and proximal tibia using a 26-G needle. Tight- and Loose-CATT were created by passing a 3-0 (0.22 mm diameter) or 4-0 (0.17 mm diameter) nylon suture, respectively, through the tunnels and tightening it to restrain ATT. The 3-0 nylon suture was thicker than the 4-0 nylon suture and exhibited superior mechanical properties (Supplementary Figure S1). In the ACL-R group, a 3-0 suture was loosely looped without restraining ATT.

### 2.3 Anterior Drawer Testing

Anterior drawer testing was performed to evaluate ATT restraint provided by the nylon suture and knee stability of the healed ACL after suture removal. We carefully removed the muscle from the knee while retaining the suture, then fixed the knee joint at 90° on the testing device (Fig. 2A). An anterior force of 0.49 N (0.05 kgf) was applied to the tibia, and radiographs were obtained before and after nylon suture removal. ATT was measured from sagittal X-ray images using ImageJ (National Institutes of Health, Bethesda, MD, USA). ATT was defined as the perpendicular distance from the most prominent point of the femoral condyle to the long axis of the tibia (Fig. 2B).

**Fig. 2.**
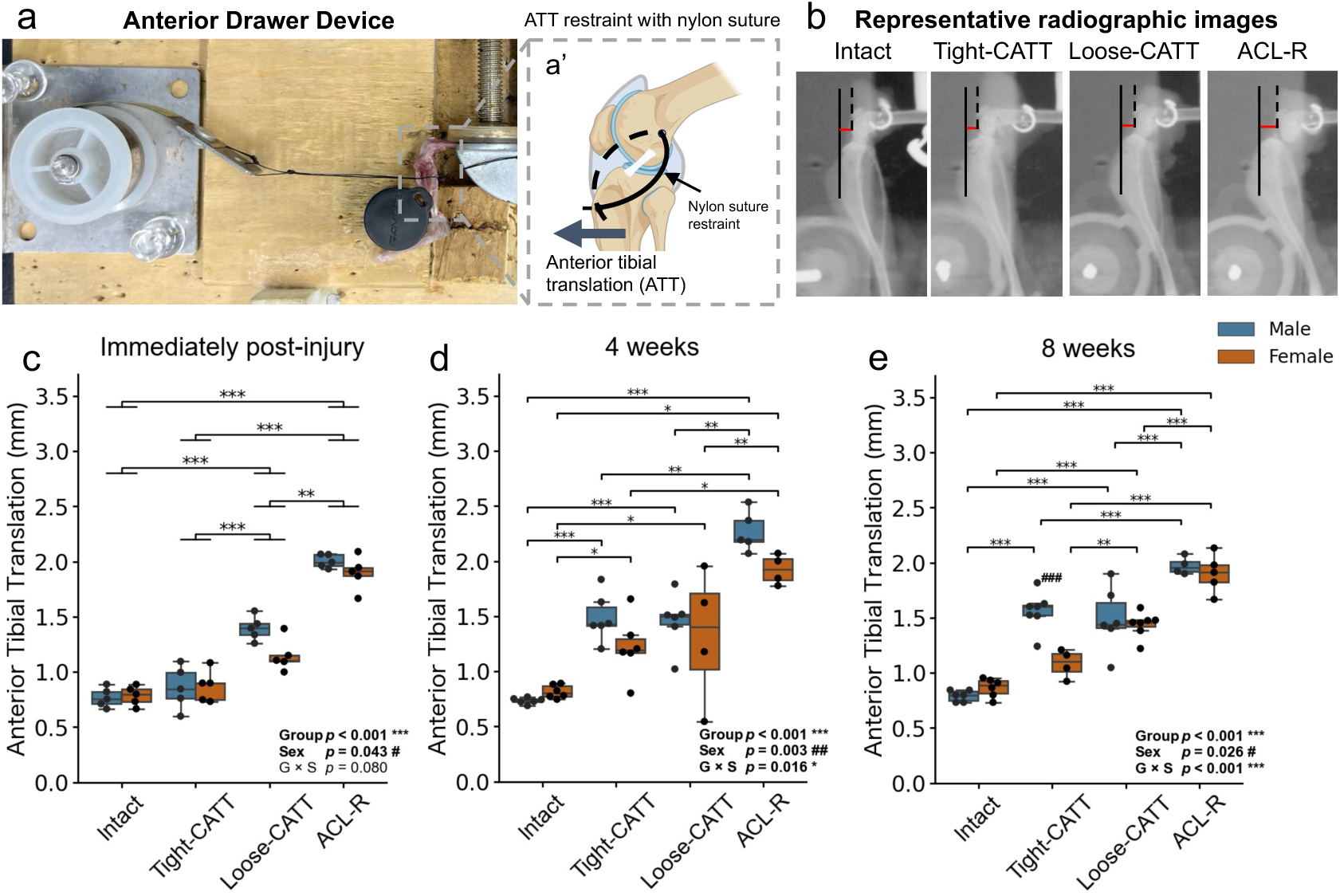
Restraint of anterior tibial translation. (A) Anterior drawer testing device with the knee fixed at 90° and (A’) schematic of ATT restraint by the nylon suture. (B) Representative X-ray images of each group. Quantification of ATT immediately after injury (C), and at 4 (D) and 8 weeks (E) after ACL injury. Box plots show individual data points (blue, male; orange, female). Asterisks indicate group differences and hash symbols sex differences. *p < 0.05, **p < 0.01, ***p < 0.001; #p < 0.05, ##p < 0.01, ###p < 0.001.

### 2.4 Histology

After anterior drawer testing, knee samples were fixed in 4% paraformaldehyde for 24–48 h, decalcified in 10% EDTA for 2 weeks, cryoprotected in sucrose, embedded in OCT, and sectioned at 12 μm. Hematoxylin and eosin (HE) staining was performed to assess ACL continuity and morphological features. ACLs showing continuous ligamentous tissue bridging the femoral and tibial remnants were classified as healed, and healing rates were calculated in each group. Picrosirius red (PsR) staining was performed to evaluate collagen organization. All samples were imaged with a fixed polarizer orientation, and the ACL long axis was consistently aligned vertically (top to bottom of the imaging field). PsR images were converted to 8-bit grayscale using Fiji (ImageJ), and a fixed threshold was applied to the ACL region of interest to quantify collagen-positive area. Fiber coherency was calculated from 10 randomly selected 150 × 150-pixel subregions using OrientationJ.

### 2.5 Immunofluorescence

Immunofluorescence staining was performed to evaluate the localization of type I collagen (COL1A1) in the healed ACL. ACL cryosections were blocked at room temperature for 1 h using DAKO Antibody Diluent with Background Reducing Components (Agilent Technologies, Inc., CA, USA). Sections were incubated overnight at 4°C with a goat polyclonal anti-COL1A1 antibody (1:200; AB758, Merck/Sigma-Aldrich, Taufkirchen, Germany), followed by incubation with an Alexa Fluor 488-conjugated anti-goat secondary antibody (1:500; ab150141, Abcam, Cambridge, UK) for 4 h at room temperature. Nuclei were counterstained with DAPI. All sections were imaged using a BZ-X710 microscope under identical imaging conditions.

### 2.6 Biomechanical Testing

Mechanical testing was performed to evaluate the failure load of the healed ACL. The harvested knee joints were dissected and frozen at −80°C. After thawing, the nylon suture and all soft tissues except the ACL were removed, and ACL healing was confirmed by microscopic observation of ligament continuity between the femoral and tibial remnants. The femur and tibia were fixed with 1.5-mm tubes to the mechanical testing machine (Univert; CellScale Biomaterials Testing, Waterloo, ON, Canada). The knee was mounted at 90° of flexion with the tensile load applied along the long axis of the ACL. After application of a 0.1-N preload, the ACL was loaded to failure at 10 mm/min. The maximum failure load and failure mode (midsubstance rupture or avulsion) were recorded.

### 2.7 Statistical Analysis

Statistical analyses were performed using Python 3.14 and R 4.5. ACL healing status was analyzed using multivariable logistic regression with Firth’s bias-reducing penalized likelihood to reduce small-sample bias. The model included group (Tight-CATT vs Loose-CATT), sex (female vs male), and assessment time point (4 vs 8 weeks) as main effects. The ACL-R group was not included in the model because no ACLs healed in this group, which precluded estimation of odds ratios. Adjusted odds ratios (ORs) and 95% confidence intervals (CIs) based on the profile penalized likelihood were reported. P values were obtained using penalized likelihood ratio tests. Continuous outcomes were assessed for normality using the Shapiro–Wilk test; because normality was not satisfied for some outcomes and subgroup sizes were small, all continuous outcomes were analyzed using an aligned rank transform (ART) (26) followed by two-way ANOVA for group, sex, and their interaction. Pairwise comparisons were performed using ART-C (27). When a significant group × sex interaction was detected, simple-effect comparisons were performed between groups within each sex and between sexes within each group. When no significant interaction was detected, pairwise comparisons were performed for factors with a significant main effect. Holm adjustment was applied to account for multiple comparisons. Cohen’s d was calculated from ART-C contrast estimates divided by the residual SD. Statistical significance was set at *p* < 0.05.

## 3 Results

### 3.1 Validation of ATT Restraint in the CATT Models

Immediately after ACL rupture, we observed no significant interaction between group and sex, but significant main effects of group (*p* < 0.001) and sex (*p* = 0.043). Immediately after injury, no significant difference was observed between the intact and Tight-CATT groups (*p* = 0.090, *d* = 0.781). These groups showed significantly lower ATT than the Loose-CATT (vs Intact: *p* < 0.001, *d* = -3.475, vs Tight-CATT: *p* < 0.001, *d* = -2.694) and ACL-R groups (vs Intact: *p* < 0.001, *d* = -5.532, vs Tight-CATT: *p* < 0.001, *d* = -4.750) (Fig. 2C). At 4 and 8 weeks, significant group × sex interactions were observed (4 weeks: *p* = 0.016, 8 weeks: *p* < 0.001). At 4 weeks, ATT in the Tight-CATT group was significantly higher than that in the intact group in both males and females (male: *p* < 0.001, *d* = 3.364, female: *p* = 0.044, *d* = 1.388), and no longer differed from that in the Loose-CATT group in either sex (male: *p* = 0.682, *d* = 0.107, female: *p* = 0.854, *d* = -0.267). No significant sex differences were observed within any group (Fig. 2D). At 8 weeks, ATT in females did not differ between the intact and Tight-CATT groups (*p* = 0.068, *d* = -1.216), whereas it was significantly higher in the male Tight-CATT group than in the intact group (*p* < 0.001, *d* = 5.041). In addition, ATT was significantly higher in males than in females within the Tight-CATT group (*p* < 0.001, *d* = 3.082). Furthermore, in females, ATT remained significantly lower in the Tight-CATT group than in the Loose-CATT group (*p* = 0.006, *d* = -1.983) (Fig. 2E).

### 3.2 ACL Healing Rate

Histologically determined ACL healing rates at 4 and 8 weeks after ACL rupture are presented in Table 1. The ACL-R group showed no ACL healing at either 4 or 8 weeks. Multivariable logistic regression analysis showed that the odds of ACL healing were significantly higher in the Tight-CATT group than in the Loose-CATT group after adjustment for sex and time point (Tight-CATT vs Loose-CATT: adjusted OR, 4.29; 95% CI, 1.26–16.90; *p* = 0.019). Female mice showed higher healing odds than male mice, although the difference did not reach statistical significance (female vs male: adjusted OR, 2.93; 95% CI, 0.85–11.72; *p* = 0.090). No significant effect of time point was observed (8w vs 4w: adjusted OR, 1.19; 95% CI, 0.34–4.18; *p* = 0.776) (Table 1).

**Table 1.** ACL healing rate. Data are presented as the number of healed and non-healed ACLs and the ACL healing rate (%). The effects of ATT restraint group, sex, and time point on ACL healing were assessed using multivariable logistic regression.

|  | 4 weeks |  |  |  | 8 weeks |  |  |  |
|  | Tight-CATT |  | Loose-CATT |  | Tight-CATT |  | Loose-CATT |  |
|  | Male | Female | Male | Female | Male | Female | Male | Female |
|---|---|---|---|---|---|---|---|---|
| Healed | 5 | 4 | 2 | 1 | 3 | 4 | 1 | 4 |
| Non-healed | 2 | 1 | 3 | 3 | 4 | 0 | 5 | 3 |
| ACL healing rate (%) | 71.4 | 80.0 | 40.0 | 25.0 | 42.9 | 100.0 | 16.7 | 57.1 |

### 3.3 Histological Evaluation of the Healed ACL

Histologically, healed ACLs in both CATT groups exhibited heterogeneous cellular organization with nuclear aggregations, unlike the more uniform appearance of the intact ACL (Fig. 3).

**Fig. 3.**
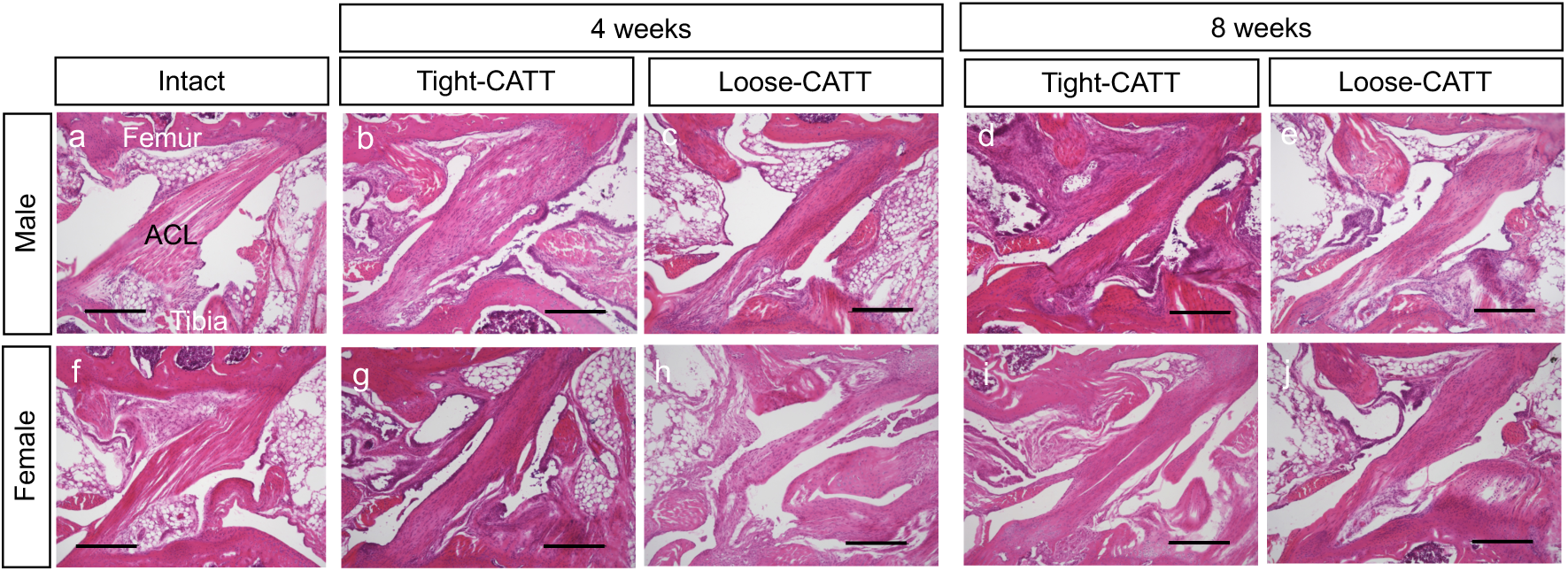
Representative HE-stained images of the healed ACL. Male (A–E) and female (F–J), Intact (A, F), Tight-CATT (B, D, G, I), and Loose-CATT (C, E, H, J) groups at 4 and 8 weeks. Scale bar = 300 μm.

Under polarized light, healed ACLs in both CATT groups exhibited greater green birefringence, indicative of thinner and less mature collagen fibers (Fig. 4A–C) and more dispersed fiber orientation (Fig. 4D–F) than the intact ACL. Quantitative analysis of the collagen-positive area revealed a significant group × sex interaction at 4 weeks (*p* = 0.009). In both males and females, the collagen-positive area was significantly lower in the Tight-CATT and Loose-CATT groups than in the intact group. In addition, the collagen-positive area was significantly higher in the Tight-CATT group than in the Loose-CATT group in females (*p* = 0.022, *d* = 2.824) (Fig. 4G). At 8 weeks, neither the group × sex interaction nor the main effect of group was significant (Fig. 4H). For fiber coherency, no significant group × sex interaction was observed, whereas the main effect of group was significant at both 4 and 8 weeks (both p < 0.001). At both 4 and 8 weeks, fiber coherency was significantly lower in the Tight-CATT and Loose-CATT groups than in the intact group. No significant differences in fiber coherency were observed between the Tight- and Loose-CATT groups at either time point (4 weeks: *p* = 0.060, *d* = 1.419; 8 weeks: *p* = 0.832, *d* = 0.146) (Fig. 4I, J).

**Fig. 4.**
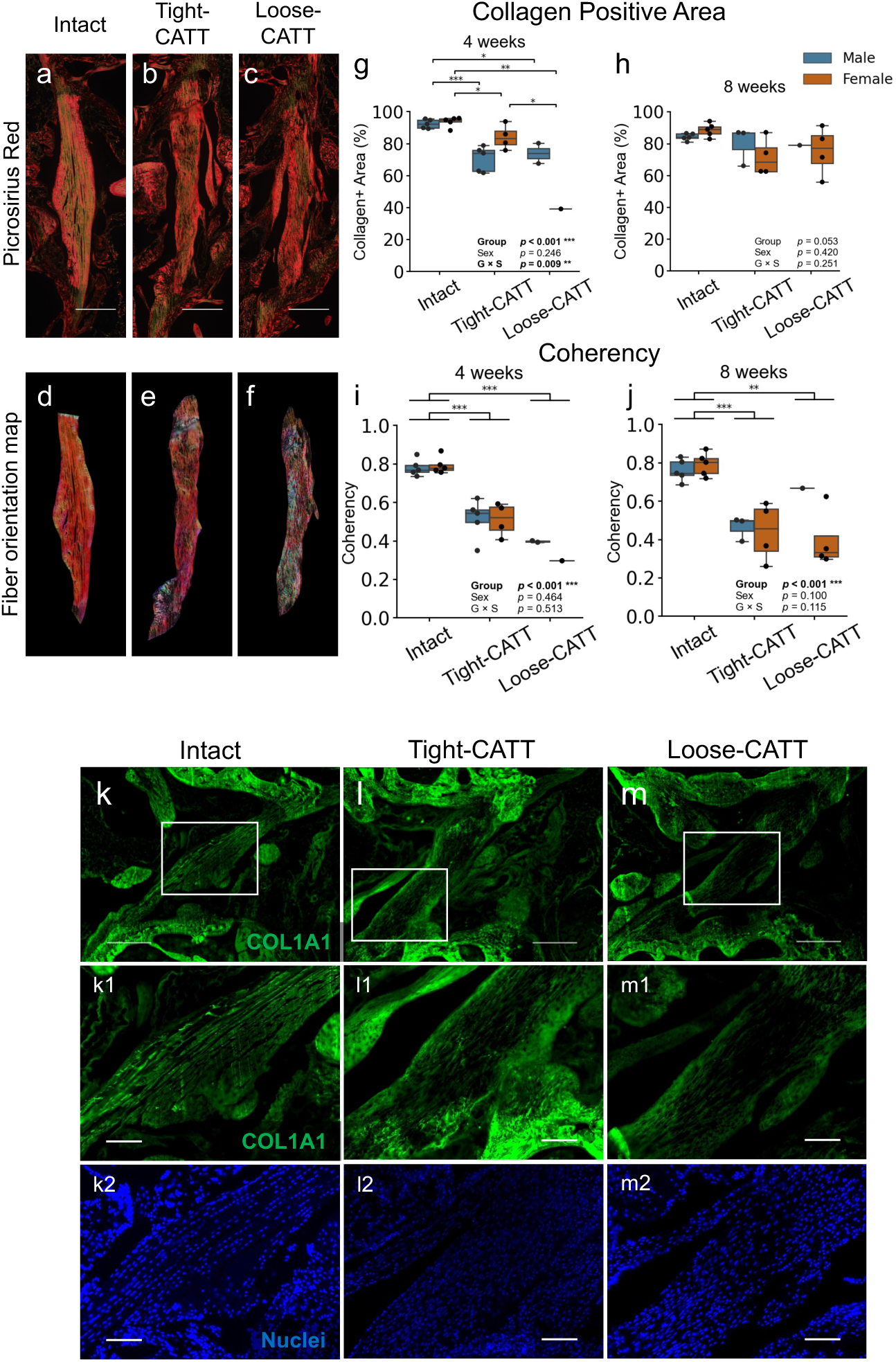
Representative PsR staining under polarized light (A–C), collagen fiber orientation maps (D–F), collagen-positive area at 4 (G) and 8 weeks (H), and fiber coherency at 4 (I) and 8 weeks (J). COL1A1 immunofluorescence is shown in K–M. A, D, K: Intact; B, E, L: Tight-CATT; C, F, M: Loose-CATT. K1–M1 and K2–M2 show magnified COL1A1 and DAPI images, respectively. Box plots show individual data points (blue, male; orange, female). Scale bars = 300 μm (A–C, K–M) and 100 μm (K1,2–M1,2). Asterisks indicate group differences. *p < 0.05, **p < 0.01, ***p < 0.001.

COL1A1 staining was relatively uniform throughout the intact ACL (Fig. 4K), whereas the healed ACLs in the Tight-CATT and Loose-CATT groups showed preferential COL1A1 localization at the ligament periphery, with relatively lower staining in the central region (Fig. 4L, M).

### 3.4 Knee Stability After ACL Healing

Significant group × sex interactions were observed at both 4 and 8 weeks (4 weeks: *p* < 0.001, 8 weeks: *p* = 0.006). At 4 weeks, ATT was significantly higher in both the Tight-CATT and Loose-CATT groups than in the intact group in males and females (Fig. 5C). At 8 weeks, ATT also remained significantly higher in both CATT groups than in the intact group in males and females (Fig. 5D). In females, ATT was significantly lower in the Tight-CATT group than in the Loose-CATT group (4 weeks: *p* < 0.001, *d* = -4.806, 8 weeks: *p* = 0.025, *d* = -1.820), whereas no significant difference was observed between the Tight-CATT and Loose-CATT groups in males (4 weeks: *p* = 0.095, *d* = 1.502, 8 weeks: *p* = 0.763, *d* = 0.352) (Fig. 5C, D). Significant sex differences were observed in all groups at 4 weeks; however, the direction of these differences was not consistent across groups (Fig. 5C). At 8 weeks, no significant sex differences were observed in any group (Fig. 5D).

**Fig. 5.**
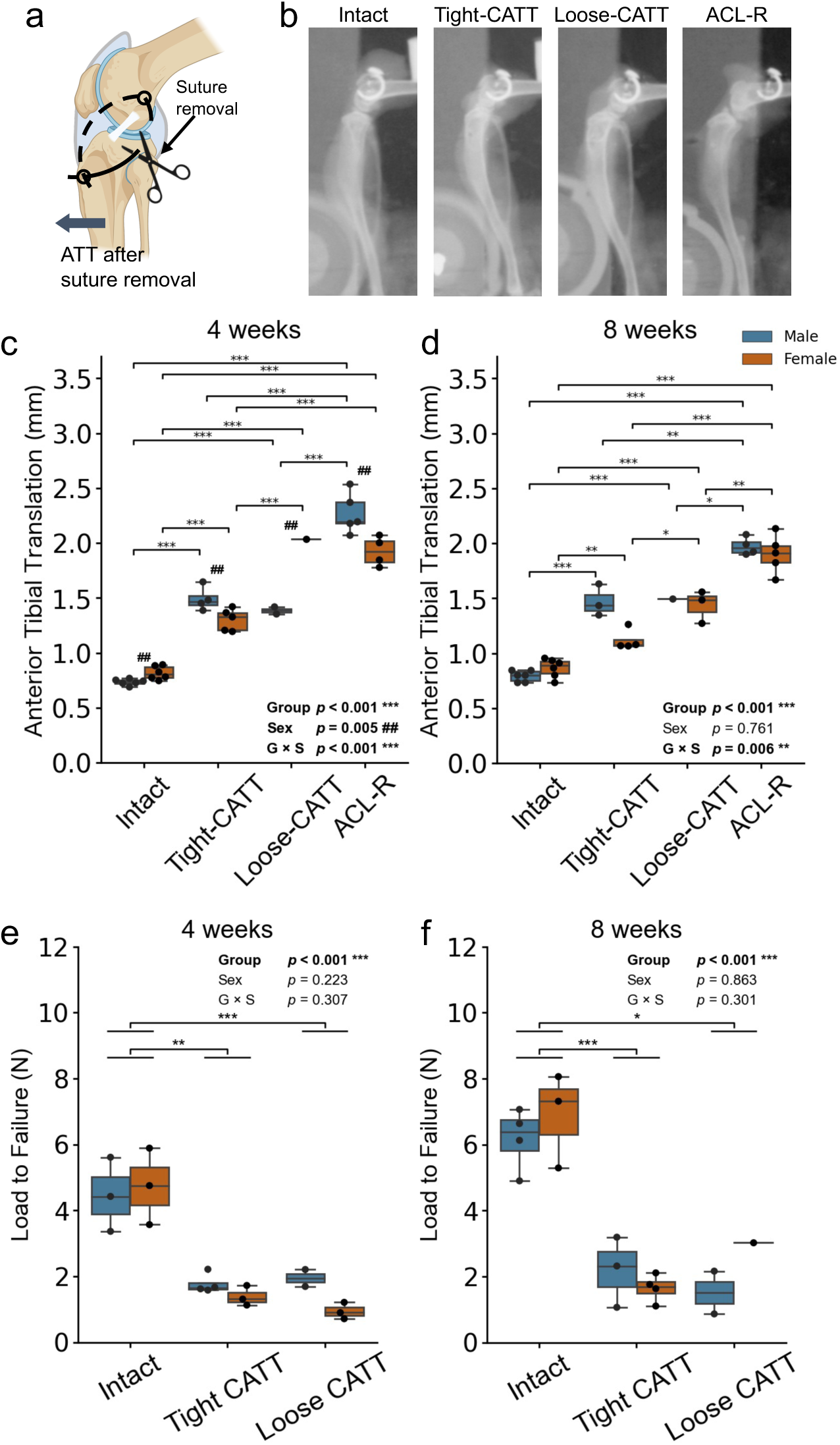
ATT measurement after removal of the nylon suture (A). Representative X-ray images of each group (B). ATT after suture removal at 4 (C) and 8 weeks (D) after ACL injury, and ACL failure load at 4 (E) and 8 weeks (F) after injury. Box plots show individual data points (blue, male; orange, female). Asterisks indicate group differences and hash symbols sex differences. *p < 0.05, **p < 0.01, ***p < 0.001; #p < 0.05, ##p < 0.01, ###p < 0.001.

### 3.5 Mechanical Properties of the ACL

All specimens ruptured at the mid-substance of the ACL. Mechanical testing showed no significant group × sex interaction at either time point, but a significant main effect of group at both 4 and 8 weeks (4 and 8 weeks: *p* < 0.001). At 4 and 8 weeks, the maximum failure load was significantly lower in the Tight-CATT and Loose-CATT groups than in the intact group. No significant differences in maximum failure load were observed between the Tight-CATT and Loose-CATT groups at either time point (4 weeks: *p* = 0.229, *d* = 0.754, 8 weeks: *p* = 0.575, *d* = - 0.417) (Fig. 5E, F). The maximum failure loads of the healed ACLs in both groups were less than half of that of the intact ACL.

## 4 Discussion

Recent clinical and preclinical studies have shown that completely ruptured ACLs can heal spontaneously under certain conditions (9–15); however, the mechanical environment required to facilitate successful ACL healing remains poorly defined. In this study, we investigated how different degrees of anterior tibial translation (ATT) restraint influence spontaneous ACL healing following injury. The most important finding was that the Tight-CATT group had significantly higher odds of ACL healing than the Loose-CATT group after adjustment for sex and time point. In contrast to our initial hypothesis, however, allowing mild residual ATT in the Loose-CATT group did not improve collagen organization or mechanical strength of the healed ACL (Fig. 4I, J, Fig. 5E, F). Taken together, these findings suggest that strict early mechanical stabilization is important for successful spontaneous ACL healing, whereas allowing mild residual ATT in the acute phase did not enhance subsequent tissue maturation or mechanical recovery.

The mechanical environment is a key determinant of ligament healing because mechanical loading can either promote or impede tissue repair. In the anterior drawer test performed with the nylon restraint maintained immediately after ACL rupture, the Tight-CATT group exhibited ATT that did not differ significantly from that of intact knees, whereas progressively greater ATT was observed in the Loose-CATT and ACL-R groups (Fig. 2C). Although ATT restraint in the Tight-CATT group had decreased to a level comparable to that in the Loose-CATT group by 4 weeks, the Tight-CATT group showed significantly higher healing odds (Table 1). This temporal relationship suggests that minimizing ATT during the early phase after ACL injury may increase the likelihood of successful bridging between the ruptured ligament remnants. Temporary extra-articular compensation for the mechanical stabilizing function of the ACL may therefore facilitate the establishment of tissue continuity during this vulnerable phase. Previous studies have shown that the injury site is filled with a mechanically fragile provisional matrix during the initial phase of ligament healing (28). Additionally, controlling abnormal joint motion suppresses the expression of matrix degradation–related factors during ligament healing (10,13,29). Thus, increased ATT during this phase may disrupt the provisional matrix and interfere with bridging between the ligament remnants, thereby reducing the likelihood of continuous ligament formation.

Beyond its effect on healing success, early ATT restraint may also influence the subsequent functional stability of the healed ACL, although this effect appeared to be sex dependent. After suture removal, ATT remained greater than that in the intact group in both CATT groups. However, among female mice, ATT was lower in the Tight-CATT group than in the Loose-CATT group (Fig. 5C, D). One possible explanation for the greater residual ATT observed in female Loose-CATT mice is elongation of the healed ligament, as progressive elongation despite restored tissue continuity has also been reported during Achilles tendon healing (30). Despite this partial improvement in functional stability, collagen fiber organization and mechanical strength of the healed ACL remained inferior to those of the intact ACL (Fig. 4I, J, Fig. 5E, F). The effects of mechanical loading on ligament healing depend on both magnitude and timing (21,31,32). Strict stabilization may therefore be most important for initial ACL healing, whereas appropriately timed loading during later stages may be required for tissue maturation and functional recovery.

Although the overall effect of sex was not statistically significant, healing rates were numerically higher in female mice in most subgroups (adjusted OR, 2.93; p = 0.090). At 4 weeks, ATT and healing rates were similar between sexes in the Tight-CATT group. At 8 weeks, however, female Tight-CATT mice showed lower ATT and a higher healing rate than male Tight-CATT mice, suggesting a possible association between maintenance of ATT restraint and successful ACL healing (Fig. 2D, E; Table 1). In the Loose-CATT group, the healing rate at 8 weeks was also numerically higher in females than in males (57.1% vs. 16.7%), despite the absence of a sex difference in ATT (Fig. 2E). Thus, measured ATT alone cannot fully explain the possible sex-related variation in healing. Sex hormones, including testosterone and estrogen, can modulate tissue repair and collagen metabolism (33–35); however, evidence that these effects influence the re-establishment of ligament continuity remains limited. Another possible explanation is a sex-related difference in spontaneous activity or cage behavior. Male mice have been reported to exhibit more frequent fighting and aggressive behaviors (36,37), which could alter joint loading and the maintenance of ATT restraint during healing. However, hormonal status, cage activity, and in vivo joint loading were not assessed in this study. Given the nonsignificant overall effect of sex and the small subgroup sizes, these observations should be considered exploratory. Whether biological sex and activity level influence spontaneous ACL healing requires further investigation.

From a clinical perspective, these findings support the concept that limiting excessive ATT during the early phase after ACL rupture may help preserve conditions favorable for tissue bridging. However, our results do not define the optimal magnitude or duration of restraint and should not be interpreted as direct evidence for a specific bracing or rehabilitation protocol. Further studies are needed to determine whether early stabilization followed by appropriately timed loading can improve both healing success and functional recovery in clinically relevant settings.

This study has several limitations. First, anatomical and biomechanical differences between murine and human knees limit direct clinical extrapolation. Second, because assessments were performed only immediately after injury and at 4 and 8 weeks, temporal changes in ATT during the early healing phase were not fully characterized. Third, the biological mechanisms underlying the association between ATT restraint and ACL healing were not directly examined. Fourth, analyses of tissue maturation, knee stability, and mechanical properties were restricted to healed ACLs. This resulted in small and unequal subgroup sizes and may have introduced selection bias, limiting both statistical power and causal interpretation of between-group differences. Finally, mice aged 8–9 weeks are not fully skeletally mature, which may limit generalization to adult patients. Further studies incorporating temporal and molecular analyses are needed to clarify the mechanobiological processes of ACL healing.

## 5 Conclusion

In summary, strict ATT restraint during the early phase was associated with a higher likelihood of spontaneous ACL healing, supporting the importance of early mechanical stabilization. Contrary to our hypothesis, allowing mild residual ATT was associated with lower healing odds and did not improve collagen organization or mechanical strength of the healed ACL. Thus, mild residual ATT during the early phase provided no apparent benefit for either ACL healing or subsequent ligament maturation, suggesting that adequate stabilization should be prioritized during the early phase of ACL healing. Clinically, these findings support further investigation of early stabilization strategies for non-operative ACL management, including the optimal magnitude and duration of restraint and the timing of subsequent loading.

## CRediT authorship contribution statement

**Riku Saito:** Conceptualization, Data curation, Formal analysis, Funding acquisition, Investigation, Methodology, Validation, Visualization, Writing – original draft, Writing – review and editing; **Kazuki Nakayama**: Formal analysis, Investigation; **Yuna Usami**: Resources; **Saaya Enomoto**: Formal analysis; **Koyo Nogi**: Validation; **Takanori Kokubun**: Conceptualization, Data curation, Funding acquisition, Methodology, Supervision, Project administration, Writing – review and editing.

## Ethics approval

All animal experiments were approved by the Animal Experiment Ethics Committee of Saitama Prefectural University (approval number: 2021-09) and conducted in accordance with the relevant institutional guidelines. The study was reported in accordance with the ARRIVE guidelines.

## Data availability

The datasets generated and/or analyzed during the current study are available from the corresponding author on reasonable request.

## Acknowledgments

We would like to thank the members of Kokubun Laboratory at Saitama Prefectural University for their cooperation.

---

**Supplementary Figure S1.**
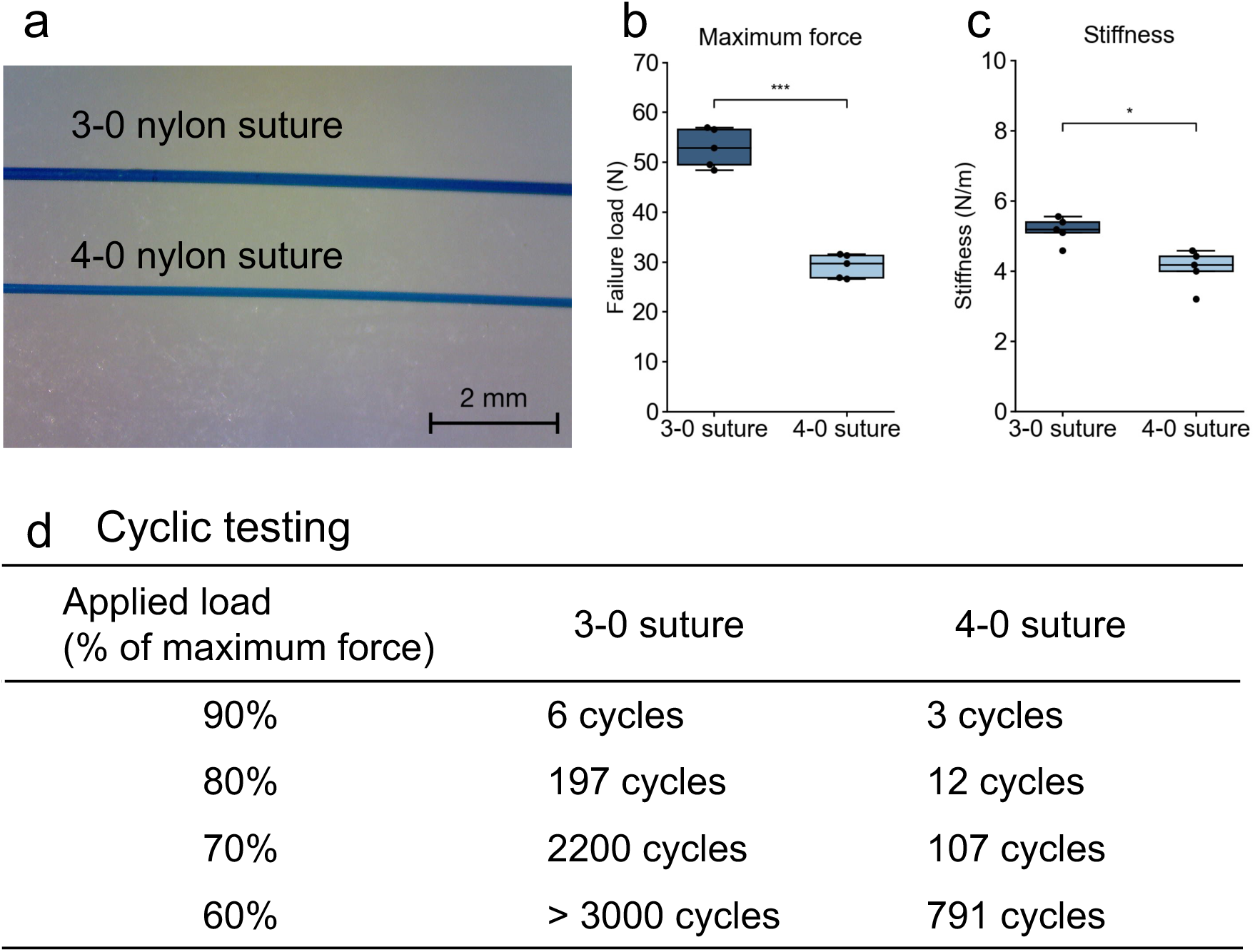
Mechanical properties of 3-0 and 4-0 nylon sutures. (a) Stereomicroscopic images of the 3-0 and 4-0 nylon sutures. Maximum failure load (b) and stiffness (c) obtained from tensile testing of each suture. (d) Failure cycles of 3-0 and 4-0 nylon sutures under cyclic loading at 90%, 80%, 70%, and 60% of the maximum force determined for each suture size. Failure cycle was defined as the number of loading cycles completed before suture failure.

## References

1. Spindler KP, Wright RW. Anterior Cruciate Ligament Tear. N Engl J Med. 2008;359(20):2135–42. doi:10.1056/NEJMcp0804745.

2. Mather RC, Koenig L, Kocher MS, Dall TM, Gallo P, Scott DJ, et al. Societal and Economic Impact of Anterior Cruciate Ligament Tears. Journal of Bone and Joint Surgery. 2013;95(19):1751–9. doi:10.2106/JBJS.L.01705.

3. Bray RC, Leonard CA, Salo PT. Correlation of healing capacity with vascular response in the anterior cruciate and medial collateral ligaments of the rabbit. J Orthop Res. 2003;21(6):1118–23. doi:10.1016/S0736-0266(03)00078-0.

4. Menetrey J, Laumonier T, Garavaglia G, Hoffmeyer P, Fritschy D, Gabbiani G, et al. α-Smooth muscle actin and TGF-β receptor I expression in the healing rabbit medial collateral and anterior cruciate ligaments. Injury. 2011;42(8):735–41. doi:10.1016/j.injury.2010.07.246.

5. Murray MM. Current status and potential for primary ACL repair. Clin Sports Med. 2009;28(1):51–61. doi:10.1016/j.csm.2008.08.005.

6. Murray MM, Fleming BC. The Biology of Anterior Cruciate Ligament Injury and Repair: Kappa Delta Ann Doner Vaughn Award Paper 2013. J Orthop Res. 2013;31(10):1501–6. doi:10.1002/jor.22420.

7. Bixby EC, Heyworth BE. Management of Anterior Cruciate Ligament Tears in Skeletally Immature Patients. Curr Rev Musculoskelet Med. 2024;17(7):258–72. doi:10.1007/s12178-024-09897-9.

8. Best MJ, Zikria BA, Wilckens JH. Anterior Cruciate Ligament Injuries in the Older Athlete. Sports Health: A Multidisciplinary Approach. 2021;13(3):285–9. doi:10.1177/1941738120953426.

9. Ihara H, Miwa M, Deya K, Torisu K. MRI of Anterior Cruciate Ligament Healing. Journal of Computer Assisted Tomography. 1996;20(2):317.

10. Kokubun T, Kanemura N, Murata K, Moriyama H, Morita S, Jinno T, et al. Effect of Changing the Joint Kinematics of Knees With a Ruptured Anterior Cruciate Ligament on the Molecular Biological Responses and Spontaneous Healing in a Rat Model. Am J Sports Med. 2016;44(11):2900–10. doi:10.1177/0363546516654687.

11. Ihara H, Kawano T. Influence of Age on Healing Capacity of Acute Tears of the Anterior Cruciate Ligament Based on Magnetic Resonance Imaging Assessment. Journal of Computer Assisted Tomography. 2017;41(2):206. doi:10.1097/RCT.0000000000000515.

12. Kano T, Kokubun T, Murata K, Oka Y, Ozone K, Arakawa K, et al. Influence of the site of injury on the spontaneous healing response in a rat model of total rupture of the anterior cruciate ligament. Connective Tissue Research. 2022;63(2):138–50. doi:10.1080/03008207.2021.1889529.

13. Morishita Y, Kanemura N, Kokubun T, Murata K, Takayanagi K. Acute molecular biological responses during spontaneous anterior cruciate ligament healing in a rat model. Sport Sci Health. 2019;15(3):659–66. doi:10.1007/s11332-019-00583-9.

14. Filbay SR, Roemer FW, Lohmander LS, Turkiewicz A, Roos EM, Frobell R, et al. Evidence of ACL healing on MRI following ACL rupture treated with rehabilitation alone may be associated with better patient-reported outcomes: a secondary analysis from the KANON trial. Br J Sports Med. 2023;57(2):91–9. doi:10.1136/bjsports-2022-105473.

15. Filbay SR, Dowsett M, Chaker Jomaa M, Rooney J, Sabharwal R, Lucas P, et al. Healing of acute anterior cruciate ligament rupture on MRI and outcomes following non-surgical management with the Cross Bracing Protocol. Br J Sports Med. 2023;57(23):1490–7. doi:10.1136/bjsports-2023-106931.

16. Butler DL, Noyes FR, Grood ES. Ligamentous restraints to anterior-posterior drawer in the human knee. A biomechanical study. J Bone Joint Surg Am. 1980;62(2):259–70.

17. Wang JHC. Mechanobiology of tendon. Journal of Biomechanics. 2006;39(9):1563–82. doi:10.1016/j.jbiomech.2005.05.011.

18. Gomez MA, Woo SLY, Amiel D, Harwood F, Kitabayashi L, Matyas JR. The effects of increased tension on healing medial collateral ligaments. Am J Sports Med. 1991;19(4):347–54. doi:10.1177/036354659101900405.

19. Andarawis-Puri N, Flatow EL. Promoting effective tendon healing and remodeling. Journal of Orthopaedic Research. 2018;36(12):3115–24. doi:10.1002/jor.24133.

20. Killian ML, Cavinatto L, Galatz LM, Thomopoulos S. The role of mechanobiology in tendon healing. Journal of Shoulder and Elbow Surgery. 2012;21(2):228–37. doi:10.1016/j.jse.2011.11.002.

21. Chen H, Li S, Xiao H, Wu B, Zhou L, Hu J, et al. Effect of Exercise Intensity on the Healing of the Bone-Tendon Interface: A Mouse Rotator Cuff Injury Model Study. Am J Sports Med. 2021;49(8):2064–73. doi:10.1177/03635465211011751.

22. Woo SLY, Vogrin TM, Abramowitch SD. Healing and Repair of Ligament Injuries in the Knee. Journal of the American Academy of Orthopaedic Surgeons. 2000;8(6):364–72. doi:10.5435/00124635-200011000-00004.

23. Provenzano PP, Martinez DA, Grindeland RE, Dwyer KW, Turner J, Vailas AC, et al. Hindlimb unloading alters ligament healing. J Appl Physiol (1985). 2003;94(1):314–24. doi:10.1152/japplphysiol.00340.2002.

24. Serdar CC, Cihan M, Yücel D, Serdar MA. Sample size, power and effect size revisited: simplified and practical approaches in pre-clinical, clinical and laboratory studies. Biochem Med (Online). 2021;31(1):27–53. doi:10.11613/BM.2021.010502.

25. Takahata K, Arakawa K, Enomoto S, Usami Y, Nogi K, Saitou R, et al. Joint instability causes catabolic enzyme production in chondrocytes prior to synovial cells in novel non-invasive ACL ruptured mouse model. Osteoarthritis and Cartilage. 2023;31(5):576–87. doi:10.1016/j.joca.2022.12.004.

26. Wobbrock JO, Findlater L, Gergle D, Higgins JJ. The aligned rank transform for nonparametric factorial analyses using only anova procedures. In: Proceedings of the SIGCHI Conference on Human Factors in Computing Systems [Internet]. New York, NY, USA: Association for Computing Machinery; 2011. p. 143–6. (CHI ’11). doi:10.1145/1978942.1978963.

27. Elkin LA, Kay M, Higgins JJ, Wobbrock JO. An Aligned Rank Transform Procedure for Multifactor Contrast Tests. In: The 34th Annual ACM Symposium on User Interface Software and Technology [Internet]. Virtual Event USA: ACM; 2021. p. 754–68. doi:10.1145/3472749.3474784.

28. Chamberlain CS, Crowley E, Vanderby R. The spatio-temporal dynamics of ligament healing. Wound Repair and Regeneration. 2009;17(2):206–15. doi:10.1111/j.1524-475X.2009.00465.x.

29. Nishikawa Y, Kokubun T, Kanemura N, Takahashi T, Matsumoto M, Maruyama H, et al. Effects of controlled abnormal joint movement on the molecular biological response in intra-articular tissues during the acute phase of anterior cruciate ligament injury in a rat model. BMC Musculoskelet Disord. 2018;19(1):175. doi:10.1186/s12891-018-2107-6.

30. Hoeffner R, Agergaard AS, Svensson RB, Cullum C, Mikkelsen RK, Konradsen L, et al. Tendon Elongation and Function After Delayed or Standard Loading of Surgically Repaired Achilles Tendon Ruptures: A Randomized Controlled Trial. Am J Sports Med. 2024;52(4):1022–31. doi:10.1177/03635465241227178.

31. Eliasson P, Andersson T, Aspenberg P. Achilles tendon healing in rats is improved by intermittent mechanical loading during the inflammatory phase: Tendon healing and early loading. J Orthop Res. 2012;30(2):274–9. doi:10.1002/jor.21511.

32. Camp CL, Lebaschi A, Cong GT, Album Z, Carballo C, Deng XH, et al. Timing of Postoperative Mechanical Loading Affects Healing Following Anterior Cruciate Ligament Reconstruction: Analysis in a Murine Model. The Journal of Bone and Joint Surgery. 2017;99(16):1382–91. doi:10.2106/JBJS.17.00133.

33. Tashjian RZ, Zitnay J, Kazmers NH, Veerabhadraiah SR, Zelada AC, Honeggar M, et al. Estrogen and testosterone supplementation improves tendon healing and functional recovery after rotator cuff repair. J Orthop Res. 2024;42(2):259–66. doi:10.1002/jor.25695.

34. Ashcroft GS, Mills SJ. Androgen receptor–mediated inhibition of cutaneous wound healing. J Clin Invest. 2002;110(5):615–24. doi:10.1172/JCI15704.

35. Sander AM, Connizzo BK. Estrogen and Progesterone Exhibit Distinct Yet Coordinated Roles in the Regulation of Tendon Extracellular Matrix Remodeling. Journal of Orthopaedic Research. 2026;44(1):e70018. doi:10.1002/jor.70018.

36. Lidster K, Owen K, Browne WJ, Prescott MJ. Cage aggression in group-housed laboratory male mice: an international data crowdsourcing project. Sci Rep. 2019;9(1):1. doi:10.1038/s41598-019-51674-z.

37. Miczek KA, Maxson SC, Fish EW, Faccidomo S. Aggressive behavioral phenotypes in mice. Behavioural Brain Research. 2001;125(1–2):167–81. doi:10.1016/S0166-4328(01)00298-4.

