## Supplemental Figure S1 for "Strict Early Restraint of Anterior Tibial Translation Favors Spontaneous Anterior Cruciate Ligament Healing: Implications for Non-Operative Management from a Murine Rupture Model"

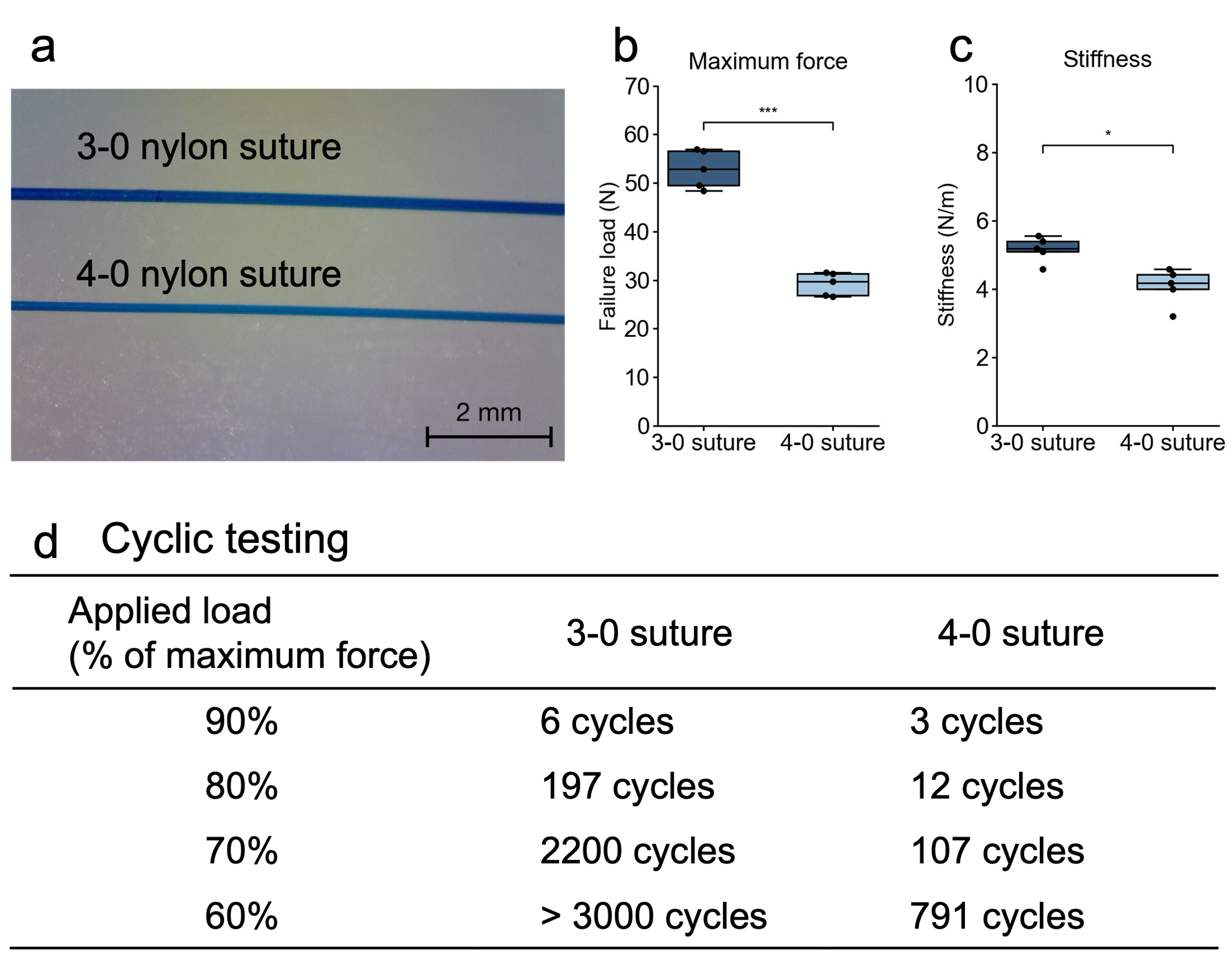

**Supplementary Figure S1.** Mechanical properties of 3-0 and 4-0 nylon sutures. (a) Stereomicroscopic images of the 3-0 and 4-0 nylon sutures. Maximum failure load (b) and stiffness (c) obtained from tensile testing of each suture. (d) Failure cycles of 3-0 and 4-0 nylon sutures under cyclic loading at 90%, 80%, 70%, and 60% of the maximum force determined for each suture size. Failure cycle was defined as the number of loading cycles completed before suture failure.
